# Exploring ID4 as a Driver of Aggression and a Therapeutic Target in Triple-Negative Breast Cancer

**DOI:** 10.1101/2025.02.07.637072

**Authors:** C Toro, S Real, S Laurito, MT Branham

## Abstract

Basal-like breast cancer (BLBC) is characterized by an aggressive clinical course, high genomic instability, and limited therapeutic options. The Inhibitor of Differentiation 4 (ID4) protein has been identified as a critical regulator of BLBC, where its overexpression correlates with poor prognosis. However, the mechanistic contributions of ID4 to BLBC tumorigenesis remain incompletely understood. In this study, we employed an integrative approach combining CRISPR-Cas9-mediated ID4 knockout, small-molecule inhibition, in vivo tumor modeling, and in silico transcriptional analyses to investigate the functional role of ID4 in BLBC.

CRISPR-Cas9-mediated knockout of ID4 in MDA-MB-231 cells resulted in significant reductions in proliferation, colony formation, and Ki67 expression, indicating a loss of aggressive phenotypic traits. In vivo xenograft studies further revealed that ID4-silenced cells exhibited markedly delayed tumor formation and a significant reduction in metastatic potential compared to controls. Kaplan-Meier survival analysis of basal-like tumors from The Cancer Genome Atlas (TCGA) dataset demonstrated that patients with low ID4 expression had improved relapse-free survival.

Gene set enrichment analysis (GSEA) of BLBC tumors stratified by ID4 expression revealed a shift toward luminal-like transcriptional programs in the ID4-low subgroup, including increased estrogen response and inflammatory signaling pathways. Furthermore, transcription factor activity analysis identified the activation of MYC, JUN, and STAT in ID4-low tumors, suggesting a transition toward a more differentiated phenotype. Finally, pharmacological inhibition of ID4 using the small-molecule degrader AGX51 effectively reduced proliferation in TNBC cells, highlighting ID4 as a potential therapeutic target.

Together, these findings establish ID4 as a key driver of BLBC aggressiveness and suggest that its inhibition may represent a viable therapeutic strategy. This study provides compelling evidence supporting the development of ID4-targeted therapies for TNBC patients, with the potential to improve clinical outcomes in this challenging disease subset.

## INTRODUCTION

Breast cancer is a highly heterogeneous disease, encompassing several subtypes with distinct molecular profiles, clinical behaviors, and therapeutic responses. Molecular classification studies have identified intrinsic subtypes, including luminal A, luminal B, HER2-enriched, basal-like, and normal-like, each associated with specific clinical outcomes^1^. Among these, basal-like breast cancer (BLBC) stands out due to its aggressive clinical course, limited therapeutic options, and poor prognosis ^2–4^. Representing a substantial proportion of triple-negative breast cancers (TNBC), BLBC is defined by the absence of hormone receptors (ER and PR) and HER2, along with the expression of basal cytokeratins (CK5/6) and epidermal growth factor receptor (EGFR) ^5,6^. These characteristics, combined with high proliferative activity and genomic instability, contribute to its propensity for recurrence and metastasis.

The lack of effective treatments for BLBC underscores the urgent need to identify key molecular drivers that could inform novel therapeutic strategies. Emerging evidence highlights the pivotal role of the Inhibitor of Differentiation (ID) protein family, particularly ID4, in this subtype. The ID protein family (ID1–ID4) comprises transcriptional regulators that lack DNA-binding domains but function as dominant-negative inhibitors of basic helix-loop-helix (bHLH) transcription factors^7^. While all ID proteins play critical roles in cell fate determination, differentiation, and proliferation, their expression and functions are context-dependent and vary across cancer types. For instance, ID1 and ID3 have been implicated in promoting angiogenesis and metastasis in various cancers^8–10^, while ID2 is often associated with cell cycle regulation ^11^. In contrast, ID4 has emerged as a unique player in BLBC, where it is frequently overexpressed and linked to poor survival outcomes ^12^. Unlike other ID proteins, ID4 is specifically implicated in maintaining stem-like properties and genomic instability in BLBC, making it a promising target for therapeutic intervention.

ID4 plays a dual role in both normal mammary gland development and the pathogenesis of BLBC ^13^. Its overexpression in BLBC is associated with impaired DNA repair proficiency and increased genomic instability, mediated through interactions with key DNA damage response proteins such as MDC1, γH2AX, and BRCA1 ^14^. These interactions highlight ID4’s critical role in driving the aggressive behavior of BLBC and further support its potential as a therapeutic target.

Despite these findings, the precise mechanisms through which ID4 drives tumorigenesis in BLBC remain incompletely understood. As a regulator of cell differentiation and proliferation, ID4 likely contributes to the stem-like properties and genomic instability characteristic of BLBC. Investigating the downstream molecular pathways regulated by ID4 may provide crucial insights into its role in promoting tumor aggressiveness and uncover potential therapeutic vulnerabilities.

This study builds upon existing knowledge by employing an integrative approach to elucidate the role of ID4 in the aggressive phenotype of BLBC. Unlike previous studies that have focused on single experimental modalities, we combine CRISPR-Cas9-mediated gene editing, small-molecule inhibition, and in silico analyses to provide a comprehensive understanding of ID4’s function. This multi-faceted approach not only allows us to validate ID4 as a critical driver of TNBC progression but also identifies actionable pathways and therapeutic vulnerabilities. By integrating these diverse methodologies, we aim to bridge the gap between mechanistic insights and translational applications, providing a foundation for the future exploration of ID4-targeted therapies.

## RESULTS

### 1. CRISPR-Cas9-Mediated ID4 Knockout Impairs Proliferation and Colony Formation in Triple-Negative Breast Cancer Cells

Given that high ID4 expression is associated with aggressive phenotypes and poor prognosis in TNBC, we sought to inhibit ID4 expression in MDA-MB-231 cell lines through CRISPR-Cas9-mediated knockout. This approach allowed for the precise targeting and disruption of the ID4 gene, enabling us to investigate its role in phenotypic traits such as cell proliferation, colony formation and migration. To ensure robust and reproducible results, we initially generated 8 individual clones and selected those that exhibited more than 80% reduction in ID4 expression, as confirmed by Western blot and qRT-PCR. This stringent selection criterion ensured that the clones used in downstream assays had consistent and significant ID4 silencing. As shown in Figure 1A, a significant reduction in ID4 mRNA and protein levels was confirmed post-knockout. Both individual clones and pools of silenced cells were utilized in downstream assays. Proliferation assays were conducted with individual clones, enabling detailed analysis of the specific effects of ID4 silencing in homogeneous populations, thus minimizing confounding factors related to cell heterogeneity. In contrast, colony formation assay was performed using pooled cells, offering a broader and more representative perspective of the impact of ID4 silencing in a heterogeneous cell population, which more closely reflects the in vivo scenario. As illustrated in Figure 1B, ID4 knockout significantly reduced proliferation and colony formation, underscoring the critical role of ID4 in these processes. To confirm reduced proliferation, we also measured Ki67 expression, a widely recognized marker of cellular proliferation, which was markedly decreased in ID4 knockout cells. Migration assays indicate reduction in migratory capacity in ID4 knockout cells, suggesting a potential role for ID4 in cell migration. While these observations are consistent with the overall loss of aggressive traits observed in ID4-silenced cells, further experiments are required to validate these findings and fully elucidate the role of ID4 in cell migration (Supplementary Figure S1).

**Figure 1.**
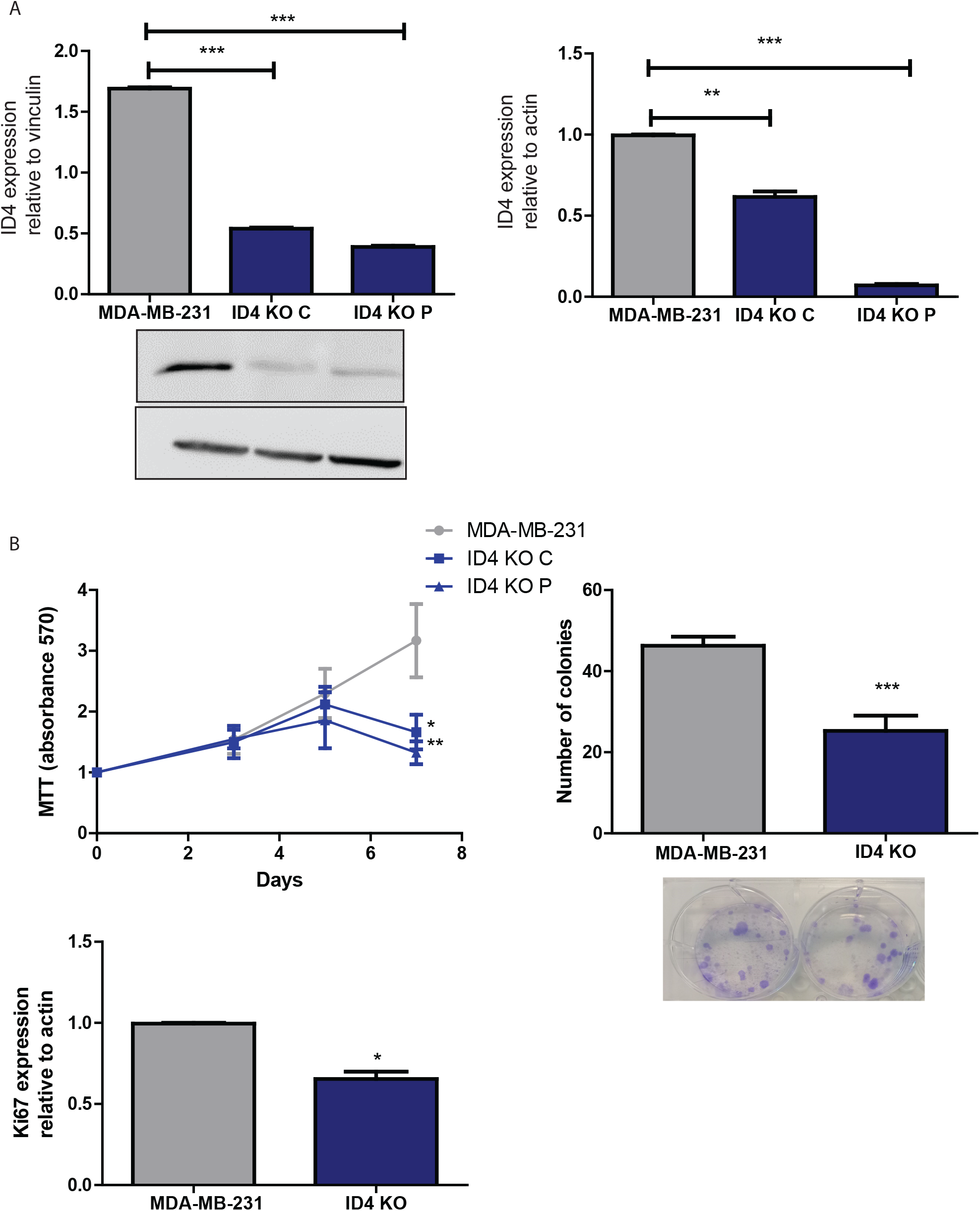
Impact of ID4 Knockout on Gene Expression and Cell Proliferation. (A) Left: Bar graph illustrating the reduction in ID4 expression following ID4 knockout using CRISPR-Cas9 in cell clones (C) and pools (P). Right: Validation of ID4 knockdown by quantitative real-time PCR. (B) Left: MTT proliferation assay demonstrating significantly reduced proliferation in ID4-silenced cells compared to controls. Right: Colony formation assay showing a marked reduction in the number of colonies formed by ID4 knockout cells. Bottom left: Bar graph indicating reduced KI67 expression in ID4 knockout cells. All graphs represent the mean ± SEM of three independent experiments. Statistical significance was assessed, and error bars indicate variability across replicates.

### 2. ID4 Silencing Suppresses Tumor Growth and Metastasis In Vivo and Correlates with Improved Patient Survival

To further assess the role of ID4 in tumor development in vivo, xenograft assays were performed using NSG immunodeficient mice. Mice were divided into two groups, with 5 mice per group: one group was inoculated with control MDA-MB-231 cells, and the other with ID4-silenced (ID4-KO) cells. Tumor growth was monitored by measuring tumor volume every three days, and growth curves were generated to compare tumor progression between the control and ID4-KO groups over time. Our results demonstrate a marked difference in tumorigenic behavior between ID4-KO and control groups. In the control group, tumor development followed the expected aggressive trajectory, necessitating humane euthanasia of the mice due to excessive tumor burden within three weeks post-inoculation. This aligns with the typical tumorigenic profile observed in this model, where even wild-type (WT) cells consistently generate detectable tumors within a similar timeframe. In contrast, ID4-KO cells exhibited a significant delay in tumor formation, with palpable tumors emerging only after approximately two months. This significant latency reflects an important reduction in the aggressiveness and tumorigenic potential of ID4-KO cells. The data suggests that ID4 plays a pivotal role in maintaining the aggressive phenotype of these cells, likely influencing key processes such as proliferation, invasion, or survival within the tumor microenvironment.

To complement this, a second silencing approach using shRNA-mediated knockdown was explored, alongside a scrambled shRNA control. As shown in Figure 2A, tumors derived from shRNA-infected cells were significantly smaller than controls (p<0.001), although the effect was less pronounced compared to CRISPR-Cas9-mediated knockout. These differences suggest that the greater efficiency, stability, and persistence of CRISPR-Cas9-mediated gene disruption surpasses the partial and potentially reversible inhibition achieved with shRNA. Notably, mice bearing ID4-silenced tumors showed no visible lung metastases, in contrast to controls (Supplementary Figure 2), highlighting the pivotal role of ID4 not only in primary tumor progression but also in metastatic dissemination.

**Figure 2.**
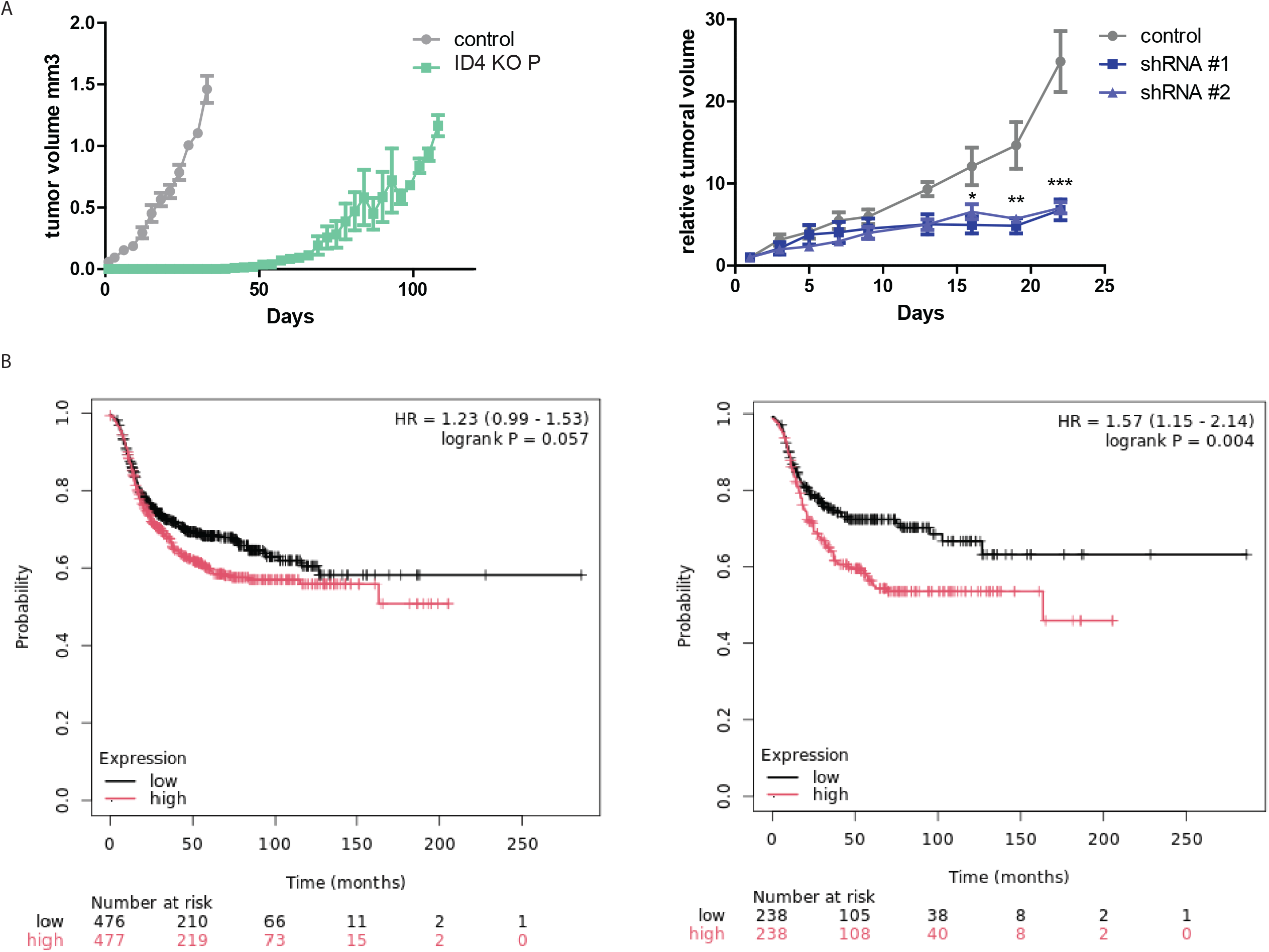
ID4 suppresses tumor development of MDA-MB-231 orthotopic xenograft in NSG mice. (A) Tumor growth curves in NSG mice xenografted with MDA-MB-231 cells. Left: ID4 knockout (CRISPR-Cas9-treated) cells did not form palpable tumors until two months post-inoculation, demonstrating a marked reduction in tumor aggressiveness. Right: Tumor volume was significantly reduced in the ID4 knockdown (shRNA-treated) group compared to the control, indicating impaired tumor progression. (B) Kaplan-Meier survival analysis of basal-like breast cancer (BLBC) patients. Left: Patients stratified by median ID4 expression levels show that low ID4 expression is associated with improved relapse-free survival (RFS). Right: Kaplan-Meier curves grouped by ID4 expression quartiles further confirm that lower ID4 expression correlates with better RFS. Hazard ratio (HR) values and statistical significance levels are indicated: *p < 0.05, **p < 0.01, ***p < 0.001. All graphs represent data from independent experiments or patient cohorts, with error bars reflecting variability. Statistical significance is annotated where applicable.

Furthermore, in silico analysis of Kaplan-Meier survival data reveals that BLBC patients with low ID4 expression exhibit better relapse-free survival (RFS) compared to those with high ID4 expression, with a p-value of 0.057 when stratified by median expression levels (Figure 2B). Notably, when ID4 expression is divided into quartiles, a more pronounced difference is observed, with patients in the first quartile (Q1) showing significantly improved RFS compared to those in the fourth quartile (Q4) (p=0.004). This observation underscores the clinical relevance of ID4, aligning with our in vivo findings that reduced ID4 expression impairs both primary tumor growth and metastasis formation. Together, these results suggest that ID4 may be a critical driver of tumor progression and a potential therapeutic target in basal-like breast cancer.

#### 3. GSEA and Transcription Factor Activity Reveal Phenotypic Shifts in Low ID4-Expressing Basal-Like Tumors Toward Luminal Characteristics

To elucidate the mechanisms underlying the reversion of aggressive phenotypic traits following ID4 knockout, we conducted in silico analyses. We selected basal-like tumors from The Cancer Genome Atlas (TCGA) dataset and stratified them into high and low ID4 expression groups based on the median ID4 expression levels. We hypothesized that the low ID4 expression group would represent conditions similar to our ID4 knockout in vitro model, thereby reflecting basal-like breast (BLB) tumors with low ID4 expression. Differential gene expression analysis and gene set enrichment analysis (GSEA) were subsequently performed to compare these groups.

Our analysis identified several significantly enriched gene sets, including hallmark interferon gamma response (NES = 3.769, p = 0.000164, FDR = 0.008190), hallmark estrogen response late (NES = 2.806, p = 0.005018, FDR = 0.095663), hallmark interferon alpha response (NES = 2.702, p = 0.006889, FDR = 0.095663), hallmark E2F targets (NES = 2.656, p = 0.007909, FDR = 0.095663), and hallmark inflammatory response (NES = 2.531, p = 0.011380, FDR = 0.095663) (Figure 3A). Among these, the hallmark estrogen response early and late gene set stood out, suggesting that estrogen signaling pathways may be active in the low ID4-expressing subgroup of basal-like breast tumors. Additionally, the significant enrichment of interferon gamma and alpha response gene sets, along with the inflammatory response gene set, suggests that tumors with low ID4 expression may exhibit increased immune and inflammatory activity.

**Figure 3:**
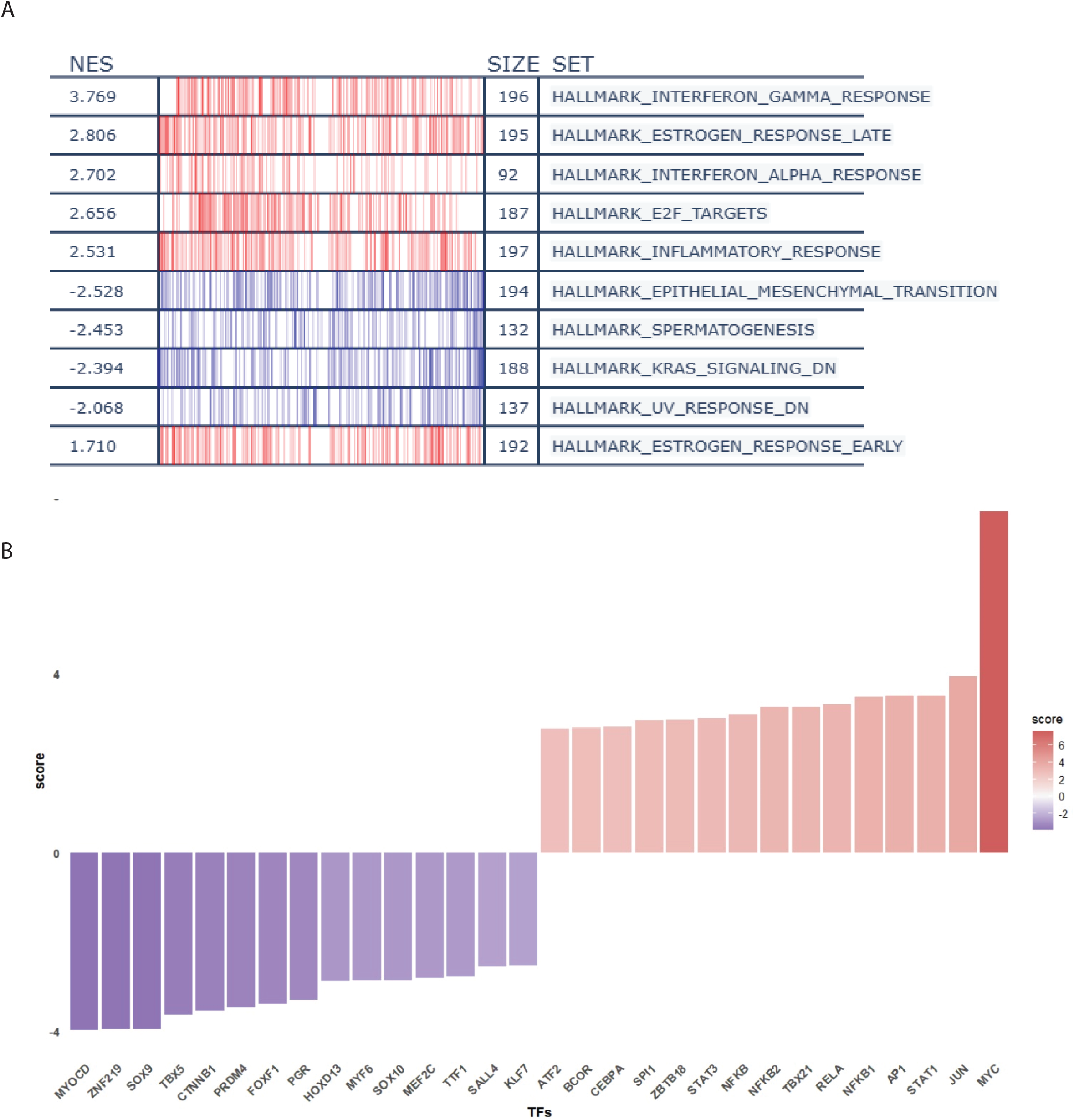
Association of Low ID4 Expression with Pathways and Transcription Factor Activity in BLBC Patients. (A) Gene Set Enrichment Analysis (GSEA) highlighting pathways associated with differentially expressed genes in patients with low ID4 expression. (B) Bar graph representing changes in transcription factor (TF) activity between patients with low and high ID4 expression in the BLBC cohort.

To further investigate the molecular mechanisms associated with ID4 expression, we performed a transcription factor (TF) activity analysis using the decoupleR package in R, comparing low and high ID4 expression groups. This analysis identified 42 TFs with increased activity and 44 TFs with decreased activity in the low ID4 group (Table 1 and Figure 3B). Among the TFs with increased activity were MYC, JUN, and STAT1, which are known to interact with estrogen receptors and are typically more active in luminal breast tumors. Conversely, TFs with decreased activity included factors such as MYOCD, SOX9, SOX10, and SALL4, which are involved in differentiation and the aggressive behavior of basal-like breast tumors.

These findings, together with our previous in silico observations, suggest that within the basal-like subtype, tumors with low ID4 expression exhibit a phenotype more closely resembling differentiated, hormone-responsive luminal tumors.

### 4. AGX51-Induced ID4 Degradation Reduces Cell Proliferation in TNBC Cells

Based on our observation that ID4 silencing induces a less aggressive phenotype in TNBC cell lines, we propose that ID4 silencing could serve as a potential therapeutic strategy for TNBC patients. To further investigate this, we examined the effects of AGX51, a recently developed small molecule inhibitor known to degrade ID proteins, including ID4 ^15,16^. MDA-MB-231 cells were treated with 40 µM AGX51 for 72 hours to evaluate its impact on cell behavior and ID4-associated pathways. First, we confirmed that AGX51 effectively induces the degradation of ID4 in TNBC cells (Figure 4 A). Following this confirmation, we assessed the impact of AGX51 on cell proliferation. As shown in Figure 4 B, treatment with AGX51 resulted in a significant reduction in cell proliferation in vitro. To further validate this effect, we assessed the expression of Ki67, a widely recognized proliferation marker in clinical practice. Consistent with the observed reductions in proliferation, Ki67 expression was significantly diminished following AGX51 treatment.

**Figure 4:**
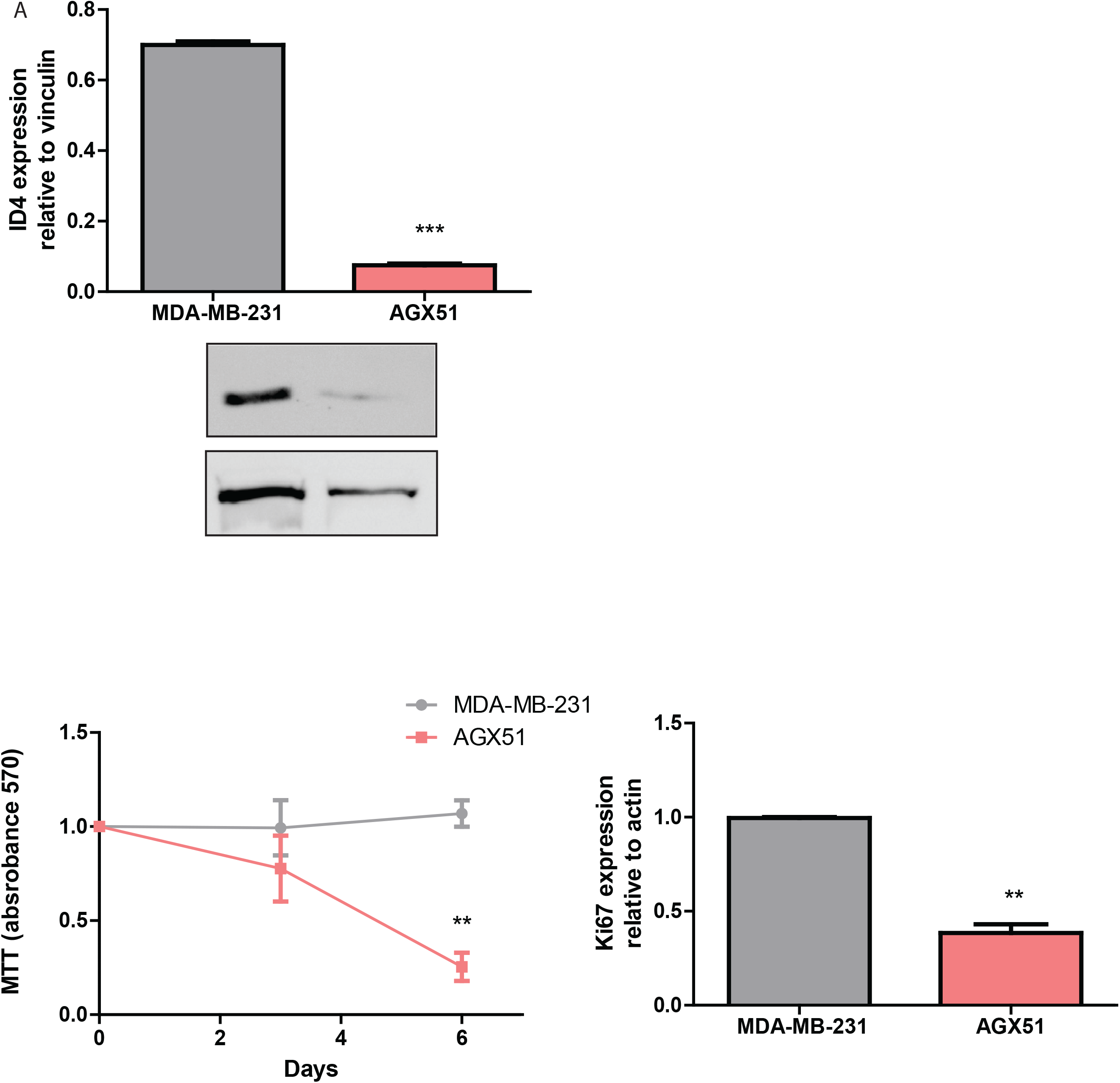
Effects of AGX51 Treatment on ID4 Expression and Cell Proliferation. (A) Bar graph showing the reduction in ID4 expression following AGX51 treatment. (B) Left: MTT proliferation assay indicating a significant decrease in cell proliferation in AGX51-treated cells compared to controls. Right: Bar graph illustrating reduced KI67 expression in AGX51-treated cells. All graphs represent the mean ± SEM of three independent experiments. Statistical significance was assessed, with error bars indicating variability across replicates. **p<0.01, ***p<0.001.

### 5. AGX51 Restores BRCA1 Expression and Disrupts Oncogenic Signaling Pathways in TNBC Cells

Given that ID4 has been proposed to inhibit BRCA1 expression ^17,18^, we evaluated BRCA1 levels following AGX51 treatment. ID4 is known to interact with DNA damage response proteins, including BRCA1, at fragile chromatin sites. Our results demonstrated a significant increase in BRCA1 expression after AGX51-induced degradation of ID4 (Figure 5 A), suggesting a potential restoration of DNA damage repair mechanisms. Although AGX51 can target other ID proteins, the observed increase in BRCA1 expression strongly implicates ID4 as a key target in this context. This finding is particularly significant in the context of TNBC, where defects in DNA repair pathways, such as homologous recombination deficiency (HRD), are common. By restoring BRCA1 expression, AGX51 may enhance the ability of TNBC cells to repair DNA damage. This dual mechanism—targeting both ID4 and restoring BRCA1 function—positions AGX51 as a promising therapeutic candidate for TNBC patients.

**Figure 5:**
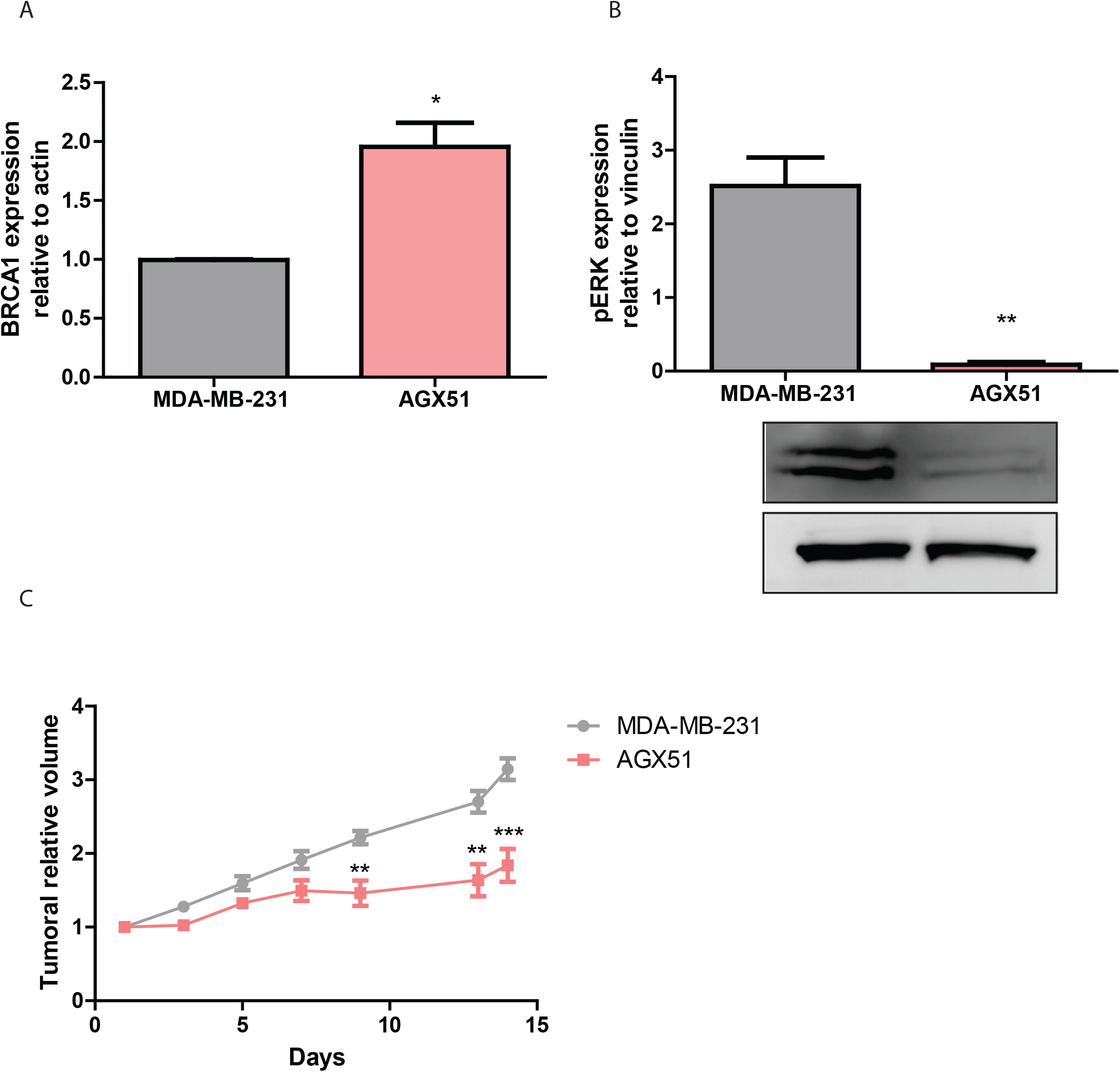
Impact of AGX51 Treatment on BRCA1 Expression, pERK Activity, and Tumor Growth in NSG Mice (A). The bar graph on the left illustrates a significant increase in BRCA1 expression following AGX51 treatment. This increase was statistically significant, with a p-value of less than 0.05, indicating a robust upregulation of BRCA1 in response to the treatment. (B). The bar graph shows a marked reduction in pERK activity in AGX51-treated cells compared to controls. The reduction in pERK activity was statistically significant, demonstrating the efficacy of AGX51 in inhibiting this pathway. (C) Tumor growth curves in NSG mice xenografted with MDA-MB-231 cells are presented, comparing control and AGX51-treated groups. The AGX51-treated group exhibited significantly slower tumor growth compared to the control group, with statistical significance. This suggests that AGX51 effectively inhibits tumor progression in vivo. All graphs represent the mean ± SEM of three independent experiments. Statistical significance was assessed using appropriate tests, and error bars indicate variability across replicates. **p<0.01, ***p<0.001.

To further elucidate the molecular effects of AGX51, we investigated its impact on key signaling pathways in MDA-MB-231 cells, focusing on pathways central to TNBC progression. Notably, AGX51 treatment led to a marked reduction in the phosphorylation levels of ERK1 and ERK2 (pERK), indicating suppressed activation of the ERK signaling pathway (Figure 5B). The ERK pathway is a critical driver of TNBC aggressiveness, promoting cell proliferation, survival, and metastasis. High ERK activity is associated with poor prognosis in TNBC, as it enhances tumor invasiveness and resistance to therapy. By suppressing ERK1/2 phosphorylation, AGX51 disrupts this oncogenic signaling cascade, potentially reducing tumor growth and metastatic potential. This finding underscores the therapeutic potential of AGX51 in targeting multiple pathways that contribute to TNBC progression, offering a multifaceted approach to combat this aggressive disease.

### 6. In Vivo Efficacy and Tolerability of AGX51 in a TN Breast Cancer Xenograft Model

To evaluate the effects of AGX51 on tumor development in vivo, we performed xenograft assays in NSG mice. Treatment began 30 days post-implantation, when tumors reached approximately 150 mm^3^, administering 15 mg/kg of AGX5. These in vivo studies are essential for validating AGX51’s efficacy in a pre-clinical environment, allowing us to assess tumor growth, potential side effects, and drug biodistribution. Our results demonstrated a significant reduction in tumor size in AGX51-treated mice compared to the control group, underscoring its therapeutic potential for TN breast cancer (Figure 5 C). Importantly, no significant differences in weight or histopathological evaluation of major organs were observed between AGX51-treated and vehicle-treated animals, indicating that AGX51 is well tolerated at this dosage.

## DISCUSSION

This study highlights the central role of ID4 in driving the aggressive behavior of TNBC cells and presents compelling evidence for its potential as a therapeutic target. Through an integrative approach combining CRISPR-Cas9 gene editing, small-molecule inhibition, and in silico analyses, we elucidated the molecular mechanisms underpinning ID4-associated phenotypes and proposed avenues for therapeutic intervention.

Our findings confirm that ID4 promotes TNBC aggressiveness, as evidenced by reduced proliferation, colony formation, and metastatic potential upon CRISPR-Cas9-mediated ID4 knockout. These observations align with clinical data associating high ID4 expression with poor outcomes in breast cancer patients. Notably, ID4 has also been implicated in promoting angiogenesis by its regulation of pro-angiogenic cytokines such as interleukin-8 (IL-8), CXCL1, and vascular endothelial growth factor (VEGF) ^19,20^. Consistent with these findings, ID4 silencing using both CRISPR-Cas9 and shRNA approaches resulted in reduced tumor growth and apparent suppression of metastasis in vivo. While our xenograft models lacked visible lung metastases, further experiments involving dedicated metastasis assays are required to validate this link. Additionally, improved relapse-free survival in patients with low ID4 expression further underscores its therapeutic relevance in TNBC.

ID4 is overexpressed in a subset of BLBC patients, where it is associated with a stem-like phenotype and poor prognosis ^12^. To investigate this, we stratified basal-like tumors in silico into high and low ID4 expression groups. High ID4 expression correlated with hallmark gene sets such as KRAS signaling and epithelial-mesenchymal transition (EMT), both of which drive aggressive behaviors like proliferation and metastasis. KRAS plays a critical role in maintaining mesenchymal features in basal-type breast cancer cells and is preferentially activated in basal-like subtypes compared to luminal types^21^. Activation of KRAS has been shown to promote the mesenchymal traits associated with metastatic behavior, underscoring its potential as a therapeutic target for basal-type breast cancer. Similarly, the activation of EMT in TNBC is associated with increased invasiveness and poor clinical outcomes.

Conversely, low ID4 expression was associated with hallmark estrogen response late, interferon response, and inflammatory gene sets, as well as reduced activity of transcription factors (e.g., SOX9 and SOX10) that govern basal-like differentiation. These findings suggest that low ID4 expression promotes a more differentiated, hormone-responsive phenotype, potentially increasing sensitivity to endocrine therapies. Our ongoing studies are exploring whether ID4-silenced tumors and cell lines exhibit enhanced responsiveness to these treatments.

Interestingly, Baker et al. reported limited gene expression changes following ID4 silencing in basal-like cell lines, with only six differentially expressed genes identified ^22^. These include SENP3-EIF4A1, ASNS, NPIPB6, and others, which may influence the phenotypic shifts observed in our study. For instance, SENP3-EIF4A1 is implicated in mRNA decay and protein de-SUMOylation, processes that could affect proteins involved in hormone signaling ^23^. Similarly, NPIPB6, part of the nuclear pore complex, may regulate transcription factors such as MYC and STAT. Further investigation into these genes is necessary to elucidate their roles in the ID4-associated phenotype.

Building on these insights, we demonstrated the therapeutic potential of AGX51, a small-molecule inhibitor of ID proteins. AGX51 effectively degraded ID4, impairing TNBC cell proliferation both in vitro and in vivo. In addition to reducing proliferation, AGX51 restored BRCA1 expression, suggesting a dual mechanism: enhancing DNA damage repair and disrupting oncogenic signaling. Furthermore, AGX51 treatment reduced ERK1/2 phosphorylation, a critical driver of TNBC aggressiveness via the MAPK/ERK pathway. High ERK activity is a poor prognostic marker in TNBC, associated with increased invasiveness and ROS production ^24^. By targeting ERK1/2, AGX51 may mitigate these traits, reducing tumor invasiveness and oxidative stress to improve therapeutic outcomes.

Lastly, the safety profile of AGX51 supports its viability as a therapeutic option. In vivo studies showed that AGX51 was well-tolerated, with no significant effects on body weight or major organ histopathology, highlighting its potential as a safe and effective treatment for TNBC.

In conclusion, this study underscores the significance of ID4 as a driver of TNBC progression and highlights AGX51 as a promising therapeutic candidate. Further studies are warranted to validate these findings and assess their clinical applicability in overcoming TNBC’s aggressive nature.

### Limitations

While our study provides compelling evidence of ID4’s role in TNBC and the therapeutic potential of AGX51, some limitations warrant consideration. First, the reliance on MDA-MB-231 cells, a single TNBC cell line, may limit the generalizability of our findings. Future studies should validate these results across additional TNBC models. Second, although AGX51 exhibits strong efficacy and tolerability in preclinical models, its off-target effects and pharmacokinetics in humans require further investigation. Lastly, the long-term impact of ID4 silencing or inhibition on tumor evolution and resistance mechanisms remains to be explored.

## METHODS

### Cell Culture

Human breast cancer cell line MDA-MB-231 was obtained from the American Type Culture Collection (ATCC, Manassas, VA, USA). Cells were cultured in Dulbecco’s Modified Eagle Medium (DMEM) (Gibco, Life Technologies, Grand Island, NY, USA) supplemented with 10% fetal bovine serum (FBS) (Serendipia Lab S.A., Hipólito Yrigoyen, CABA, Argentina), 100 U/mL penicillin, and 100 μg/mL streptomycin (Gibco). Cultures were maintained at 37 °C in a humidified atmosphere containing 5% CO_2_.

### ID4 Knockout and Knockdown

#### _CRISPR-Cas9 Knockout

ID4 knockout was performed using a CRISPR-Cas9 system (pRP[CRISPR]-EGFP/Puro-hCas9-U6>rID4) obtained from VectorBuilder. The plasmid targeting ID4 was transfected into MDA-MB-231 cells using polyethylenimine (PEI) as the transfection reagent. Post-transfection, cells were subjected to puromycin selection (2 µg/mL) for 3 days to ensure stable integration of the construct. ID4 knockout efficiency was validated using Western blotting and quantitative PCR (qPCR).

#### _shRNA Knockdown

For ID4 knockdown, shRNA constructs were designed, cloned into a lentiviral vector, and packaged into lentiviral particles by co-transfecting HEK293T cells with the shRNA plasmid, packaging plasmid, and envelope plasmid. Viral supernatants were filtered and used to transduce MDA-MB-231 cells. Transduced cells underwent puromycin selection (2µg/mL) for 3 days. Knockdown efficiency was confirmed by qPCR and Western blot analysis.

### Gene Expression Analysis by Real-Time PCR (qPCR)

Total RNA was extracted using TriPure reagent (Roche). RNA concentration and purity were determined using a NanoDrop spectrophotometer (Labocon). One microgram of RNA was reverse-transcribed to cDNA using M-MLV Reverse Transcriptase (INBIO Highway). cDNA was diluted to 4 ng/μL and stored at -20 °C. Real-time PCR was conducted with 16 ng cDNA per reaction using the Master qPCR SYBR Green Kit (INBIO Highway). Primer sequences used were: ID4 forward: 5’-TCCCGCCCAACAAGAAAGTC-3’, ID4 reverse: 5’-CCAGGATGTAGTCGATAACGT-3’, BRCA1 forward: 5’-TGCTCTTCGCGTTGAAGAAGT-3’, BRCA1 reverse: 5’-TGATCAACTCCAGACAGATGGG-3’, KI67: forward 5’-TGACCCTGATGAGAAAGCTCAA-3’ and reverse 5’-CCCTGAGCAACACTGTCTTTT-3’ and ACTB (β-actin) forward: 5’-TGACGTGGACATCCGCAAAG-3’, ACTB reverse: 5’-CTGGAAGGTGGACAGCGAGG-3’. The analysis was performed using AriaMx Software version

2.0 (Agilent, Santa Clara, CA, USA). Relative expression normalization of genes of interest was carried out using β-actin gene expression as endogenous reference control by the ΔΔCq method.

### Protein Expression Analysis by Western Blotting

MDA-MB-231 cells were lysed in buffer (0.5% Triton X-100, 150 mM NaCl, 5 mM EDTA, 1 M Tris–HCl, pH 7.5) containing a protease inhibitor cocktail (Halt™, Thermo Fisher Scientific). Protein concentrations were measured using the Pierce™ BCA Protein Assay Kit (Thermo Fisher Scientific). Equal amounts (30µg) of protein were separated on 12% SDS-PAGE gels and transferred to nitrocellulose membranes. Membranes were blocked in 5% non-fat dry milk in PBS, incubated overnight at 4 °C with primary antibodies: Anti-ID4 (1:1000, Cusabio, #PA002998), Anti-pERK (1:500, Cusabio #RA013456A185phHU), Anti-vinculin (1:5000, Sigma-Aldrich).

After washing, membranes were incubated with horseradish peroxidase-conjugated secondary antibodies (anti-mouse, 1:10,000; anti-rabbit, 1:3000, Invitrogen) for 1h at room temperature. Protein bands were detected using enhanced chemiluminescence (ECL, BPS Bioscience), visualized using a LAS Fujifilm 4000 imaging system (GE Healthcare Life Sciences), and quantified with ImageJ 1.53K software.

### In vitro AGX51 Treatment

MDA-MB-231 cells were seeded at a density of 1 × 10LJ cells/well in 6-well plates and allowed to adhere overnight. AGX51 was prepared as a 20 mM stock solution in 100% ethanol and added to the culture medium at a final concentration of 40 µM. Control cells received an equivalent volume of ethanol. Treatments were conducted for 72 h unless otherwise stated.

### Proliferation Assay

Cell viability was assessed via the MTT assay. Briefly, 10 µL of 5 mg/mL MTT solution in PBS was added to each well and incubated for 2.5 h at 37 °C. Formazan crystals were dissolved in 100 µL DMSO, and absorbance was measured at 570 nm using a microplate reader.

### Colony Formation Assay

Single-cell suspensions were prepared by resuspending cells in complete growth medium at 500 cells/mL. Cells were seeded in 6-well plates (1000 cells/well) and incubated for 3 weeks at 37 °C, replacing the medium every 3–4 days. Colonies were fixed with an acetic acid and methanol solution (1:7) for 25 minutes, stained with 0.1% crystal violet for 45 minutes, and washed with PBS. Colonies (>50 cells) were counted using the Image J 1.53K software.

### Migration Assay

Cell migration was assessed using a wound-healing assay. Control and ID4-KO cells were seeded in a monolayer and allowed to reach confluence. A standardized scratch was created using a sterile pipette tip, and cells were then cultured in serum-free medium to minimize proliferation effects. Images of the wound area were captured every 24 hours using an inverted microscope. The extent of wound closure was quantified using Fiji (ImageJ) software by measuring the remaining wound area at each time point.

### Bioinformatic Analysis

Gene set enrichment analysis (GSEA) was performed using the UCSC Xena Browser (https://xenabrowser.net) accessed between July and December 2024). Basal-like breast cancer data were divided into low and high ID4 expression groups based on median expression. The MSigDB_Hallmark gene set library was used to identify enriched pathways. Only tumors with an ESTIMATE score indicating at least 70% purity were included in the analysis.

Transcription factor Activity analysis. The decoupleR package was employed to infer transcription factor (TF) activities from gene expression data. RNA sequencing data from The Cancer Genome Atlas (TCGA) database were used, focusing on Basal-like breast cancer patients. Patients were stratified into low and high ID4 expression groups based on the median ID4 expression levels. Gene expression data were normalized using the Trimmed Mean of M-values (TMM) method to account for differences in library sizes.

Differentially expressed genes (DEGs) were identified using the DESeq2 package in R, with a false discovery rate (FDR) < 0.05 and a log2 fold change > 1 as the cutoff criteria. The decoupleR package integrates multiple statistical approaches to extract biological signatures from prior knowledge, emphasizing gene sets regulated by transcription factors. Gene-level statistics, such as t-values obtained from DESeq2, were used as input for the enrichment analysis.

TF activity scores were calculated using the decoupleR function, leveraging gene expression data and curated prior knowledge databases. This approach enabled the identification of key transcription factors associated with differential ID4 expression.

### Kaplan-Meier Plotter Database Analysis

Survival analysis was conducted using the Kaplan-Meier Plotter database (http://kmplot.com/analysis). Breast cancer data, stratified by basal-like subtype, were analyzed with the JetSet best probe set. ID4 expression was categorized based on median or quartile values.

### Xenograft Generation

Highly immunosuppressed NOD SCID Gamma (NSG) mice (Jackson Laboratory, RRID:IMSR_JAX:005557) were used. Six-week-old female mice (20 g) were anesthetized with 4% isoflurane in oxygen and injected with 1 × 10□ MDA-MB-231 cells (control and ID4-silenced) into the 4th mammary fat pad. For AGX51 treatment, tumors were allowed to develop before initiating treatment. Once tumors reached a volume of 150mm^3^, AGX51 was administered at a dose of 15 mg/kg every other day for 12 days. Tumor volumes were calculated as: Volume (mm^3^) = (Length × Width^2^) / 2. Mice were sacrificed using a CO_2_ chamber, and tumors were excised for analysis. All animal procedures were conducted in compliance with ethical guidelines and were approved by the Institutional Animal Care and Use Committee (CICUAL) of the National University of Cuyo, Mendoza, Argentina (Aval Nº 233/2023.)

### Statistical Analysis

GraphPad Prism v5.03 was used for all statistical analyses and graph generation.

### Article drafting

We used ChatGPT, an AI-powered language model developed by OpenAI, solely for language-related assistance in the composition of this research paper, with no influence on the content or research outcomes

## AUTHOR CONTRIBUTIONS

CT and SR performed the experiments. SRL and MTB performed in silico analysis. MTB conceived the study, designed the research, and wrote the manuscript. All authors read and approved the final manuscript.

## FUNDING

This work was supported in part by a grant from the National Agency of Scientific and Technological Promotion (FONCyT-ANPCyT) under grant number PICT 2018-N° 01573. Additional support was provided by the Convocatoria a Proyectos de Investigación de Cátedras 2022-2023 from the Facultad de Ciencias Médicas, Universidad Nacional de Cuyo, under Resolución Decano N° 25/2022.

## DATA AVAILABILITY

The gene expression data used in this study were obtained from The Cancer Genome Atlas (TCGA), which is freely available at: https://xenabrowser.net/. No specific permissions are required to access this dataset. Additionally, survival analysis was performed using the Kaplan-Meier Plotter, a publicly available tool accessible at: http://kmplot.com/.

## Figure Legends

Supplementary Figure S1. ID4 knockout reduces migratory capacity in MDA-MB-231 cells. Representative images of the scratch migration assay at T0 (0 hours) and T48 (48 hours) for MDA-MB-231 cells and ID4 knockout (ID4 KO) cells. The lower panel shows a bar chart quantifying wound closure (%), indicating reduced migration in ID4 KO cells compared to controls. Data are presented as mean ± SEM from two independent experiments (*n* = 2). Statistical significance was determined using Student’s *t*-test.

Supplementary Figure S2. Tumor Growth and Lung Metastasis in Control and ID4-Silenced Xenograft Models. (A) Representative images comparing tumor size in control and ID4-silenced groups. The visual comparison highlights the significant reduction in tumor growth in the ID4-silenced group. (B) Lung metastasis images from control and ID4-silenced mice. Arrows indicate the presence of metastatic nodules in the control group, which are absent in the ID4-silenced group. These findings underscore the role of ID4 in promoting tumor growth and metastatic dissemination.

## Notes

### Competing Interest Statement

The authors have declared no competing interest.

