## Supplementary figures and images for "Exploring ID4 as a Driver of Aggression and a Therapeutic Target in Triple-Negative Breast Cancer"

### Supplementary Figure 1

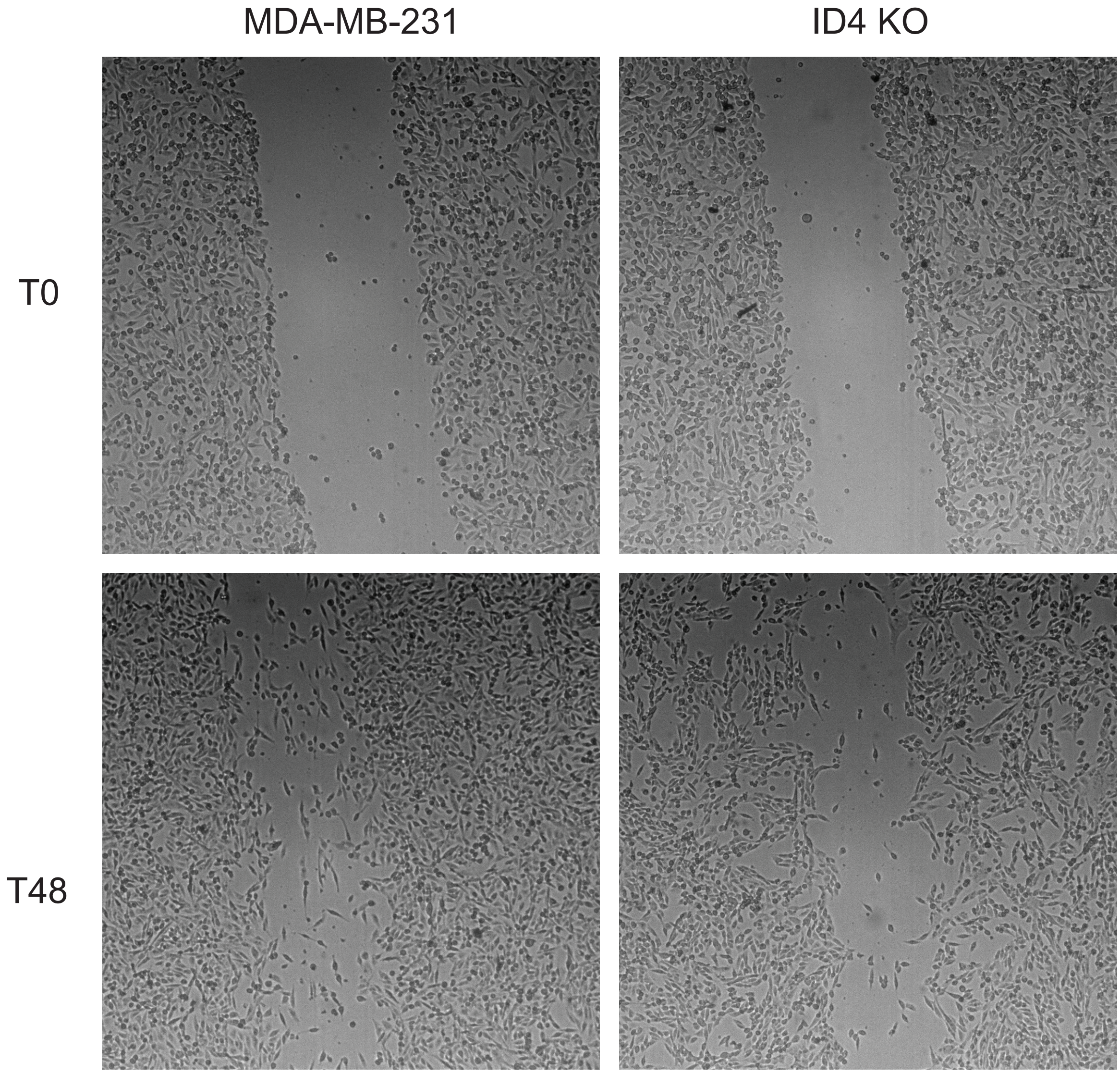

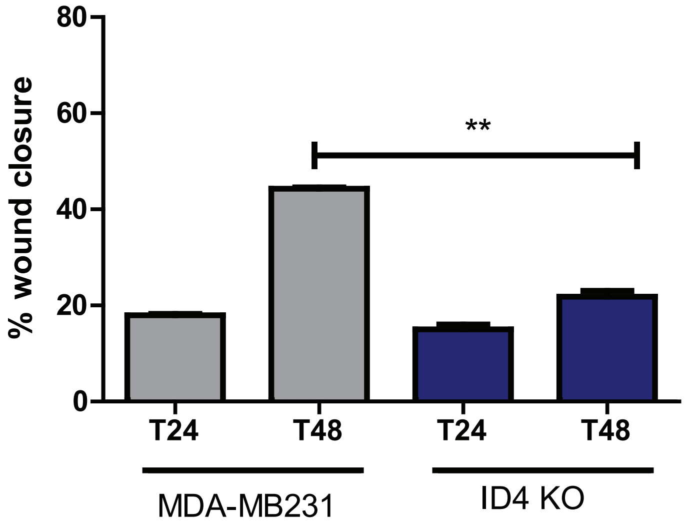

### Supplementary Figure 2

MDA-MB-231 ID4 silenced (shRNA)

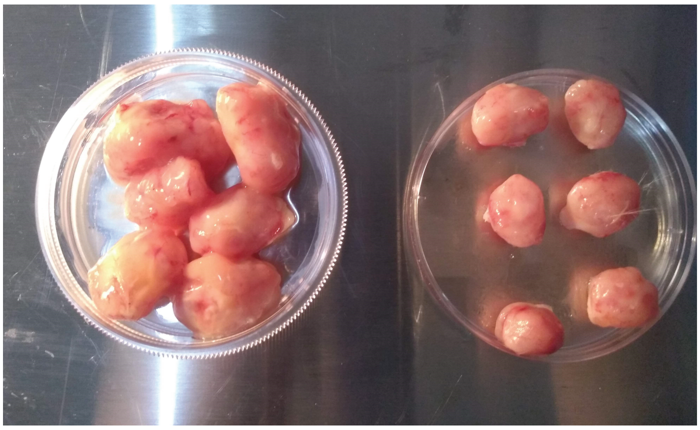

MDA-MB-231 ID4 silenced (shRNA)

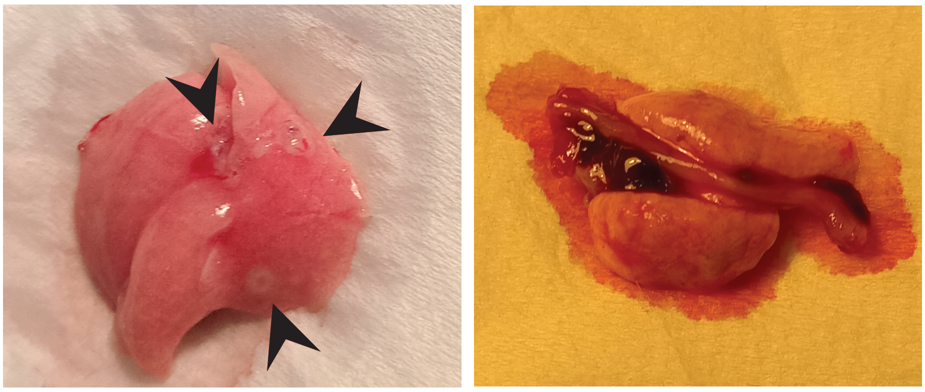
