## Supplementary material for "Exploring ID4 as a Driver of Aggression and a Therapeutic Target in Triple-Negative Breast Cancer": Table 1

**Table 1: Expression and Significance of TF**

| **TF** | **Score** | **p-value** |
| --- | --- | --- |
| MYC | 7.631 | 2.47E-14 |
| JUN | 3.932 | 8.48E-05 |
| STAT1 | 3.522 | 0.00043 |
| AP1 | 3.511 | 0.00045 |
| NFKB1 | 3.489 | 0.00049 |
| RELA | 3.31 | 0.00094 |
| TBX21 | 3.247 | 0.00117 |
| NFKB2 | 3.247 | 0.00117 |
| NFKB | 3.106 | 0.0019 |
| STAT3 | 3 | 0.00271 |
| ZBTB18 | 2.968 | 0.003 |
| SPI1 | 2.958 | 0.0031 |
| CEBPA | 2.803 | 0.00507 |
| BCOR | 2.793 | 0.00523 |
| ATF2 | 2.77 | 0.00561 |
| CEBPE | 2.732 | 0.0063 |
| HDAC7 | 2.663 | 0.00775 |
| STAT5A | 2.653 | 0.00798 |
| IRF3 | 2.599 | 0.00937 |
| BRD4 | 2.515 | 0.01192 |
| RELB | 2.51 | 0.01208 |
| HIF1A | 2.465 | 0.01369 |
| ETV5 | 2.415 | 0.01573 |
| HDAC5 | 2.395 | 0.01662 |
| IRF1 | 2.358 | 0.01838 |
| TBP | 2.298 | 0.02158 |
| ZFPM2 | 2.289 | 0.0221 |
| SPIC | 2.271 | 0.02316 |
| HR | 2.259 | 0.02391 |
| ZBED1 | 2.235 | 0.02545 |
| CEBPG | 2.203 | 0.02764 |
| NR3C1 | 2.172 | 0.02986 |
| SRSF2 | 2.158 | 0.03091 |
| REL | 2.149 | 0.03165 |
| THRB | 2.09 | 0.03662 |
| APEX1 | 2.089 | 0.0367 |
| HSF1 | 2.047 | 0.04066 |
| ZNF143 | 2.024 | 0.04297 |
| EP300 | 2.01 | 0.04445 |
| ONECUT1 | 2.005 | 0.04496 |
| HOXA2 | 2.003 | 0.04521 |
| MYT1 | 1.963 | 0.04962 |
| TOX3 | -1.979 | 0.04779 |
| LYL1 | -2.006 | 0.04486 |
| DLX5 | -2.015 | 0.04392 |
| MYOD1 | -2.018 | 0.04359 |
| PROX1 | -2.036 | 0.04179 |
| POU4F1 | -2.076 | 0.0379 |
| CXXC1 | -2.087 | 0.03694 |
| KLF13 | -2.091 | 0.03652 |
| GLI1 | -2.108 | 0.03502 |
| NKX3-1 | -2.108 | 0.03501 |
| GATA4 | -2.167 | 0.03026 |
| NEUROG3 | -2.192 | 0.02843 |
| SRF | -2.202 | 0.02767 |
| TBX6 | -2.21 | 0.02712 |
| HOXB9 | -2.212 | 0.02695 |
| ZNF263 | -2.221 | 0.0264 |
| SALL1 | -2.221 | 0.02637 |
| GATA5 | -2.256 | 0.02406 |
| MYF5 | -2.271 | 0.02317 |
| DLX2 | -2.274 | 0.023 |
| ELK3 | -2.278 | 0.02277 |
| HDAC9 | -2.284 | 0.0224 |
| HOXC6 | -2.302 | 0.02135 |
| FOXP2 | -2.364 | 0.01808 |
| NCOA2 | -2.389 | 0.01688 |
| PLAGL1 | -2.397 | 0.01655 |
| MEIS2 | -2.407 | 0.01611 |
| MYOG | -2.418 | 0.01561 |
| HOXB2 | -2.475 | 0.01332 |
| KLF7 | -2.539 | 0.01112 |
| SALL4 | -2.549 | 0.01081 |
| TTF1 | -2.761 | 0.00577 |
| MEF2C | -2.812 | 0.00493 |
| SOX10 | -2.854 | 0.00432 |
| MYF6 | -2.859 | 0.00425 |
| HOXD13 | -2.879 | 0.00399 |
| PGR | -3.305 | 0.00095 |
| FOXF1 | -3.381 | 0.00072 |
| PRDM4 | -3.468 | 0.00053 |
| CTNNB1 | -3.536 | 0.00041 |
| TBX5 | -3.632 | 0.00028 |
| SOX9 | -3.95 | 7.85E-05 |
| ZNF219 | -3.954 | 7.73E-05 |
| MYOCD | -3.977 | 7.00E-05 |
